# Noradrenaline and ATP regulate white adipocyte adiponectin exocytosis: disturbed adrenergic and purinergic signalling in obesity-associated diabetes

**DOI:** 10.1101/2021.08.19.456508

**Authors:** Saliha Musovic, Ali M. Komai, Marina Kalds Said, Yanling Wu, Ingrid Wernstedt Asterholm, Charlotta S. Olofsson

## Abstract

White adipocyte adiponectin exocytosis is triggered by cAMP and a concomitant increase of cytosolic Ca^2+^ potentiates its release. White adipose tissue is richly innervated by sympathetic nerves co-releasing noradrenaline (NA) and ATP that may act on receptors in the adipocyte plasma membrane to increase cAMP via adrenergic receptors and Ca^2+^ via purinergic receptors, respectively. Here we determine the importance of NA and ATP for the regulation of white adipocyte adiponectin exocytosis, at the cellular and molecular level, and we specifically detail the ATP signalling pathway. Immunohistochemical staining demonstrates that tyrosine hydroxylase (enzyme involved in catecholamine synthesis) is dramatically reduced in inguinal white adipose tissue (IWAT) isolated from mice with diet-induced obesity; this is associated with diminished levels of NA in IWAT and with lowered serum adiponectin. Adiponectin exocytosis (measured as increase in plasma membrane capacitance and as secreted product) is triggered by NA or ATP alone in cultured and primary mouse IWAT adipocytes, and enhanced by a combination of the two secretagogues. The ATP-induced adiponectin exocytosis is largely Ca^2+^-dependent and activated via P2Y2 receptors (P2Y2Rs) and the Gq11/PLC pathway. Adiponectin release induced by the nucleotide is abrogated in adipocytes isolated from obese/diabetic mice and this is associated with ∼70% reduced abundance of P2Y2Rs. The NA-triggered adiponectin exocytosis is likewise abolished in “obese adipocytes”, concomitant with a 50% lower gene expression of beta 3 adrenergic receptors (β_3_ARs). The NA-stimulated adiponectin secretion does not contain Ca^2+^-dependent components. Collectively, our data suggest that sympathetic innervation is a principal regulator of adiponectin exocytosis and that disruptions of this control are associated with the obesity-associated reduction of circulating levels of HMW adiponectin.

**Key point list:**

- White adipose tissue is richly innervated by sympathetic nerves that co-release noradrenaline (NA) and ATP.
- Protein levels of tyrosine hydroxylase and NA are dramatically decreased in white adipose tissue from obese/diabetic mice, concomitant with reduced serum levels of high-molecular weight (HMW) adiponectin.
- NA and ATP stimulate white adipocyte adiponectin exocytosis via beta adrenergic and purinergic receptors respectively. The ATP-induced adiponectin secretion is chiefly Ca^2+^-dependent and activated via the P2Y2/Gq11/PLC pathway.
- The purinergic signalling is abrogated in adipocytes from obese/diabetic mice, due to reduced abundance of P2Y2Rs. The response to NA is likewise abolished in “obese adipocytes”, associated with lowered gene expression of beta 3 adrenergic receptors (β_3_ARs).
- We propose that sympathetic innervation is central in regulation of adiponectin exocytosis via co-secretion of NA and ATP and that this control is disrupted in obesity-associated diabetes, leading to lower circulating levels of HMW adiponectin.

## Introduction

Adiponectin, the most abundant peptide hormone secreted by white adipocytes, has favourable metabolic actions in whole body energy homeostasis (Ruan and Dong, 2016; Wang and Scherer, 2016). Individuals with obesity-associated diabetes typically display hypoadiponectinemia and preserved adiponectin levels protect against development of metabolic disease (Kadowaki et al., 2006; Spranger et al., 2003). Although adiponectin was first described more than 20 years ago (Scherer et al., 1995) and the awareness of the endocrine function of adipose tissue originates from around the same time (Kershaw and Flier, 2004), the regulation of adiponectin secretion remains inadequately investigated. Our own previous work demonstrates that adiponectin is secreted via cAMP-triggered exocytosis of adiponectin-containing vesicles, and activation of Epac1 (*E*xchange *P*rotein directly *A*ctivated by *c*AMP, isoform 1; Holz et al., 2006). The cAMP-stimulated exocytosis is Ca^2+^-independent but can be potently augmented by an elevation of intracellular Ca^2+^ ([Ca^2+^]_i_). Ca^2+^ is also essential for the maintenance of adiponectin secretion over prolonged time-periods (adiponectin vesicle replenishment; El Hachmane et al., 2015; Komai et al., 2014). In later work, we defined adrenergic signalling as a physiological trigger of adiponectin exocytosis, via activation of beta 3 adrenergic receptors (β_3_ARs; Komai et al., 2016).

White adipose tissue is abundantly innervated by sympathetic nerves that co-release noradrenaline (NA) and ATP (Bartness et al., 2010). The finding that adiponectin exocytosis is triggered by catecholamines (Komai *et al*., 2016) proposes that the sympathetic nervous system (SNS) controls adiponectin secretion and suggests a possible regulatory role also for sympathetically released ATP. In addition to its function as an intracellular energy source, ATP has extensive roles as an extracellular signalling molecule acting on purinergic receptors in more or less all known cell types, including endocrine cells (Burnstock, 2014). Here we demonstrate a disruption of sympathetic innervation and reduced noradrenaline levels in adipose tissue isolated from mice with diet-induced obesity and that this is associated with lower circulating levels of the high-molecular weight (HMW) form of adiponectin. We therefore determine how extracellularly applied NA and ATP affect adiponectin exocytosis/secretion at a cellular and molecular level, *in vitro* and *ex vivo* using adipocytes isolated from metabolic healthy or obese/diabetic mice. We specifically detail the ATP signalling pathway involved in the stimulated adiponectin release.

## Methods

### Ethical Approval

All animal work was approved by the Regional Ethical Review Board in Gothenburg and experimental work was performed in agreement with guidelines.

### Cell and animal work

Mature 3T3-L1 adipocytes (ZenBio) and adipocytes from inguinal white adipose tissue (IWAT) isolated from 10-20 weeks old male C57BL/6J mice were used. Cells were differentiated and isolated as previously described (Komai *et al*., 2014; Komai *et al*., 2016) using reagents from Life Technologies or Sigma-Aldrich. Mice (5-weeks old) were fed chow (Global Diet #2016, Harlan-Teklad) or high fat diet (HFD; 60% kcal from fat; D12492, Research Diets Inc.) during 8 weeks. Mice were housed in groups of 10 with unlimited access to water and food and maintained on a 12-hour dark/light cycle. Mice were anesthetized with Isoflurane (2-4%, 0,2L/min). Unconsciousness was confirmed when the animals did not show any pain reflex when pinching paws and tail. Thereafter, the animal was terminated with CO_2_ or decapitation. For adiponectin secretion measurements, cells were incubated for 30 or 60 minutes at 32°C, as described (Komai *et al*., 2016). The primary adipocytes were diluted to 10-15% volume/volume. The extracellular solution (EC) contained (in mM): 140 NaCl, 3.6 KCl, 2 NaHCO_3_, 0.5 NaH_2_PO_4_, 0.5 MgSO_4_, 5 HEPES (pH 7.4 with NaOH), 2.6 CaCl_2_ and 5 glucose. Test substances were added as indicated. EC aliquots and cell homogenates were stored at −80 °C.

### Blood glucose, serum insulin and adiponectin levels

Mice were fasted during 4 hours. Trunk blood was collected during the termination. Blood glucose and serum insulin concentrations were measured using a glucose meter (Bayer Contour XT) and a mouse insulin ELISA kit (No 10-1247-01; Mercodia), respectively. Total and high-molecular weight adiponectin was measured by ELISA (EZMADP-60K; EMD Millipore and MBS028367; MyBiosource).

### Electrophysiology and [Ca^2+^]_i_ imaging

3T3-L1 adipocytes were cultured in glass (IBL) or plastic (Nunc) 35mm Petri dishes. During the experiments, the cells were superfused with EC. Exocytosis (vesicle fusion with the plasma membrane) was measured as increases in membrane capacitance (Lindau and Neher, 1988) in the standard whole-cell configuration of the patch-clamp technique (Komai *et al*., 2014). Cells were clamped at −70 mV. The pipette-filling solutions consisted of (in mM): 125 K-glutamate, 10 KCl, 10 NaCl, 1 MgCl_2_, 3 Mg-ATP and 5 HEPES (pH 7.15 with KOH). The solution was supplemented with (in mM): IC-1: 10 BAPTA; IC-2: 0.1 cAMP and 0.05 EGTA; IC-3: 0.1 cAMP and 10 EGTA; IC-4: 9 CaCl_2_, 0.1cAMP and 10 EGTA.

Intracellular Ca^2+^ concentrations ([Ca^2+^]_i_) were recorded with dual-wavelength ratio imaging in cells loaded with Fura-2 AM (Life Technologies), as previously described (Astrom-Olsson et al., 2012). Excitation wavelengths were 340 and 380 nm and emitted light was collected above 510 nm.

Measurements were carried out at 32°C.

### Quantitative real-time RT-PCR

Total RNA, extracted and purified using TRIzol (Life Technologies) and ReliaPrep™ RNA Cell Miniprep System (Promega), was reverse transcribed to cDNA using qScript Flex cDNA Kit (Quanta Biosciences). The SYBR Select Master Mix (Life Technologies) was used for quantitative RT-PCR. Primer sequences and gene symbols are shown in Supplementary Table 1.

### ELISA work

Secreted adiponectin was measured with mouse ELISA DuoSets (R&D Systems) and expressed in relation to total protein content (Bradford protein assay). Intracellular cAMP levels were measured in cell homogenates using Cyclic AMP XP Assay Kit (No. 4339; Cell Signalling). For measurements of NA levels, 10-50 mg tissue was homogenized (1 mM EDTA and 4 mM sodiummetabisulfite, pH 7.4) using a TissueLyser II (Qiagen). Samples where adjusted to the same weight/volume percentage by adding a volume of homogenization buffer and the NA content was measured by ELISA (BA E-5200, Labor Diagnostika Nord, Nordhorn, Germany) according to the manufacturer’s protocol. P2Y2R protein levels were determined in cell lysate with specific mouse ELISA (catalog # MBS7200782, MyBioSource).

### Immunohistochemical staining

Paraffin sections were incubated with Tyrosine Hydroxylase antibody (1∶300, BioSite Cat#LS-B3443) at 4°C overnight and thereafter with a secondary biotinylated antibody for 1 hour at room temperature. Sections were developed using diaminobenzidine, counterstained with haematoxylin and viewed by conventional light microscopy (Olympus BX60, Japan). Quantification was done using ImageJ.

### Data analysis

The rate of capacitance increase (ΔC/Δt) was measured at indicated time-points by application of linear fits and using OriginPro (OriginLab Corporation), as described (Komai *et al*., 2014). The free [Ca^2+^] in pipette solutions was calculated as defined in (Komai *et al*., 2014). For [Ca^2+^]_i_ imaging, the absolute [Ca^2+^]_i_ was calculated using equation 5 of (Grynkiewicz et al., 1985); Kd=224nM).

### Statistics

The statistical significance of variance between two means was calculated using Student’s t-test, (paired or unpaired) and ANOVA was used to determine statistical significance between two or more groups, and/or conditions. All data are presented as mean values ±SEM for designated number of experiments. Individual data points are typically shown. Statistics were performed with GraphPad.

## Results

### Disturbed sympathetic innervation and reduced noradrenaline levels in adipose tissue from mice with diet-induced obesity

The importance of sympathetic nervous system (SNS) innervation for white adipose tissue lipolysis has been investigated in some detail (Bartness et al., 2014). Obesity-induced reduced sympathetic sensitivity of adipose tissue has also been suggested to underlie dysregulation of leptin production and perhaps synthesis or secretion of other adipose tissue secretory products (Rayner, 2001). Subcutaneous adipose tissue has been proposed to be the most important adipose tissue depot for the control of circulating adiponectin levels (Lihn et al., 2004; Meyer et al., 2013) and our own previous work on adiponectin exocytosis has mainly focused on mouse subcutaneous inguinal white adipose tissue (IWAT) and human abdominal subcutaneous adipocytes (Brannmark et al., 2020; El Hachmane *et al*., 2015; Komai *et al*., 2014; Komai *et al*., 2016). We therefore examined the SNS innervation in IWAT isolated from mice fed chow or high-fat diet (HFD) through 8 weeks, by immunohistochemical staining for tyrosine hydroxylase (TH). TH is the rate-limiting enzyme for catecholamine biosynthesis and a sympathetic nerve marker. As visualised in Fig. 1A and quantified in Fig. 1B, TH was abundant in IWAT from lean animals whereas the signal was strikingly reduced in fat tissue isolated from the obese/diabetic mice. Moreover, NA levels were markedly lower in IWAT from HFD-fed mice compared to adipose tissue from chow-fed animals (Fig. 1C). Measurements of serum adiponectin levels in lean and obese mice (8 weeks of chow or HFD diet), demonstrated a >40% reduction of the high-molecular weight (HMW) to total adiponectin ratio in HFD-fed mice (Fig. 1D), thus similar to what we have previously reported (Komai *et al*., 2016). Our results indicate that a disturbance of adipose tissue SNS innervation exists in obesity-associated diabetes. The fact that sympathetic nerves co-release NA and ATP (Bartness *et al*., 2010), that increases adipocyte intracellular cAMP and Ca^2+^ respectively (El Hachmane et al., 2018; Kelly et al., 1989; Laplante et al., 2010; Lee et al., 2005), suggests an important role of the SNS for regulation of adiponectin secretion. Moreover, the disturbed SNS innervation in adipose tissue from obese and diabetic mice together with the reduced HMW/total adiponectin, indicate a connection between defect innervation and obesity-associated hypoadiponectinemia (Kadowaki *et al*., 2006; Spranger *et al*., 2003). Applying a combination of electrophysiological recordings and biochemical measurements, we aimed to define how NA and ATP affects adiponectin exocytosis at a cellular and molecular level. *In vitro/ex vivo* effects of the catecholamine and the nucleotide were thus studied in cultured adipocytes and in primary adipocytes isolated from lean and obese/diabetic mice.

**Fig. 1.**
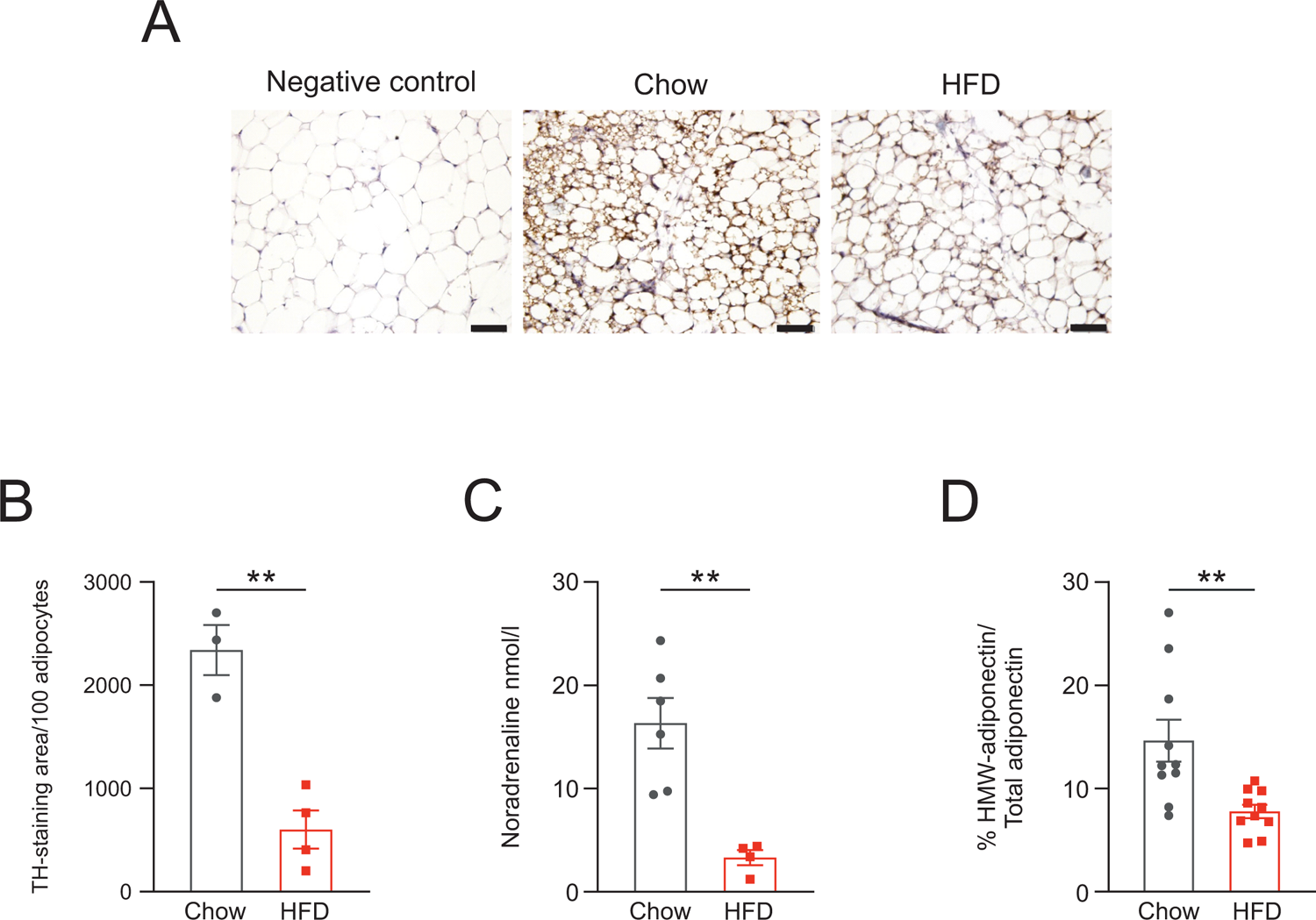
**Sympathetic innervation of adipose tissue in lean and obese mice. *A***, Representative images and (***B***) quantification of tyrosine hydroxylase (TH) staining of IWAT from chow- and HFD-fed mice (n=4 for HFD and n=3 for chow). Scale bar: 50 μm. In order to control for obesity-induced increase of adipocyte size, quantification is expressed as the stained area divided by 100 adipocytes (um^2^/100 adipocytes). ***C,*** NA content in IWAT from chow-(n=6) and HFD-fed (n=4) mice. ***D,*** Percentage HMW of total adiponectin in serum from 10 chow- and 10 HFD-fed mice. *\*P<0.05*; *\*\*P<0.01*

### Noradrenaline and ATP stimulate white adipocyte adiponectin exocytosis

IWAT adipocytes isolated from lean mice were incubated in the presence of different concentrations of ATP, alone or in combination with NA (100 nM; 30 min static incubations). Adiponectin secretion was stimulated by 100 μM ATP whereas lower concentrations of the nucleotide (0.5 and 10 μM) were without effect on adiponectin release (Fig. 2A). In agreement with data using adrenaline (ADR; Komai *et al*., 2016), NA triggered adiponectin secretion. Lower concentrations of ATP (0.5 and 10 μM) were unable to affect adiponectin release when added together with NA. The combination of NA and 100 μM ATP resulted in potent adiponectin secretion, of a magnitude significantly higher than that in the presence of ATP alone (Fig. 2A; P=0.2 *vs.* NA only). Based on those results, a concentration of 100 μM ATP was utilised throughout this study.

**Fig. 2.**
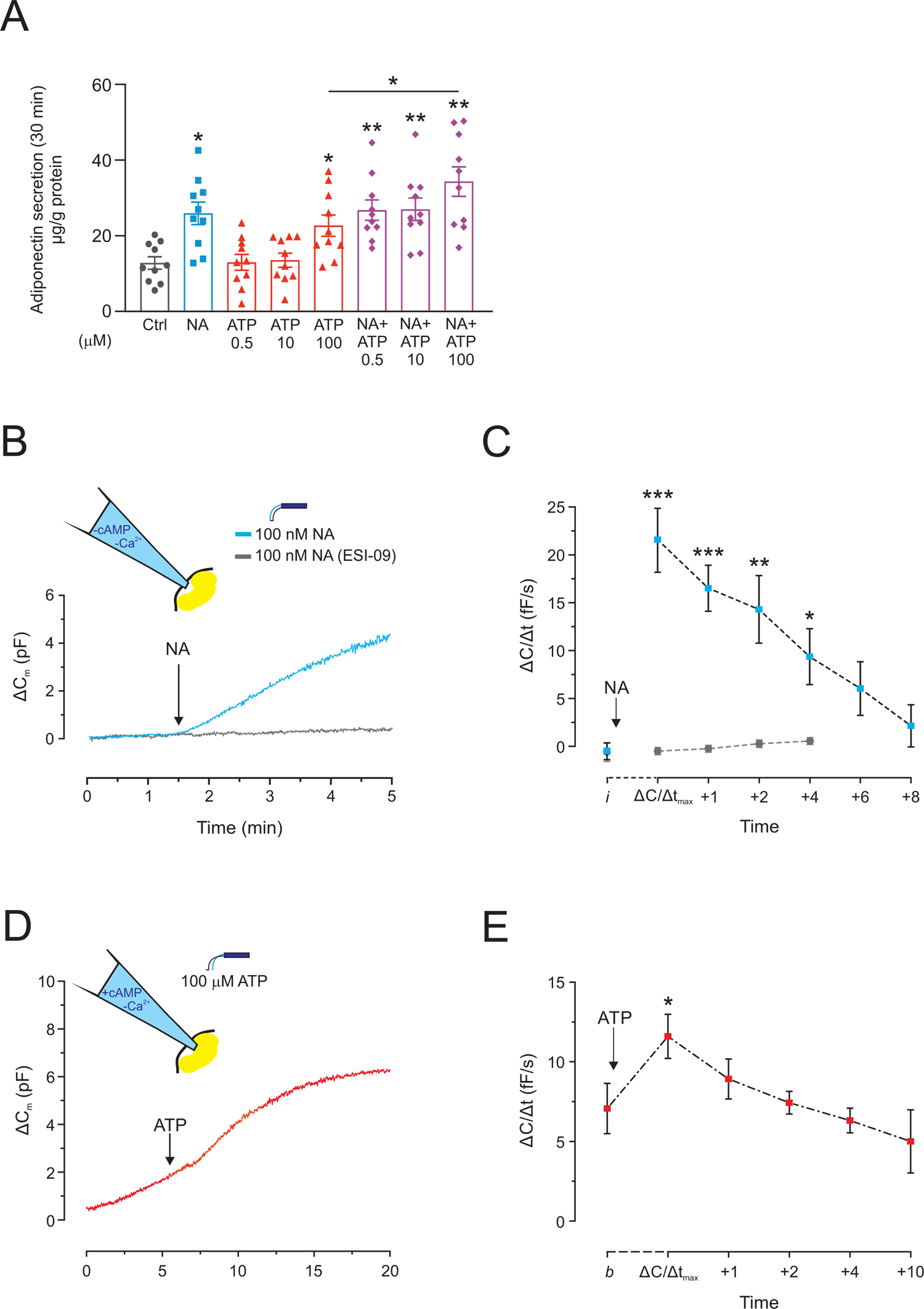
Effects of noradrenaline and extracellular ATP on white adipocyte adiponectin secretion/exocytosis. *A*, Adiponectin release in primary mouse inguinal (IWAT) adipocytes during 30 min incubation with NA and/or different concentrations of ATP, as indicated. ***B***, Two representative traces of Δ*C*_m_ for cells dialyzed with a pipette solution lacking cAMP and with NA added extracellularly (at the arrow) in the absence or presence of the Epac inhibitor ESI-09. ***C***, Average exocytotic rate (Δ*C*/Δ*t*) analysed at an indicated time point before and several time points after addition of NA in presence or absence of ESI-09. The achieved maximal exocytotic rate (Δ*C*/Δ*t*_max_) was measured and *“i”* indicates the *i*nitial rate, prior to application of NA. The indicated significance applies to the same time-point in the two traces. ***D***, Typical capacitance recording of ΔC_m_ for cells dialyzed with a Ca^2+^-depleted pipette solution containing 0.1 mM cAMP and 50 µM EGTA (IC-2), to allow for Ca^2+^ fluctuations. External ATP was added where indicated by arrow. ***E***, Average exocytotic rate (Δ*C*/Δ*t*) analysed at indicated time points after the addition of ATP. The achieved maximal exocytotic rate (Δ*C*/Δ*t*_max_) was measured and *“b”* indicates the time point just *b*efore application of ATP. Results in ***A*** are from 10 experiments with IWAT adipocytes isolated from 5 chow mice. *\*P<0.05*; *\*\*P<0.01* vs. control. Data are from 9 and 6 (ESI-09) recordings in ***C*** and 5 recordings in ***E***. *\*P<0.05*; *\*\*P<0.01; ***P<0.01*.

To investigate NA- and/or ATP-triggered adiponectin secretion in detail, we performed whole-cell capacitance measurements of exocytosis (Lindau and Neher, 1988) using cultured 3T3-L1 adipocytes, a proven relevant cell model for studying adiponectin secretion (El Hachmane *et al*., 2015; Komai *et al*., 2014; Komai *et al*., 2016). Membrane capacitance recordings allow real-time measurements of vesicular exocytosis, triggered by agents added to the pipette solution (intracellular solution; IC) or applied to the solution superfusing the cell culture dish (extracellular solution; EC). In keeping with our own previous work (El Hachmane *et al*., 2015; Komai *et al*., 2014; Komai *et al*., 2016), cells were clamped at −70 mV to circumvent activation of voltage-gated Ca^2+^ channels, shown to be expressed in adipocytes (Fedorenko et al., 2020). Cells were infused with a pipette solution lacking cAMP and Ca^2+^ (IC-1). In agreement with previous work (El Hachmane *et al*., 2015; Komai *et al*., 2014; Komai *et al*., 2016) the adipocyte membrane capacitance was unaffected by IC-1 alone. However, extracellularly applied NA triggered exocytosis and the stimulatory effect was abolished in cells pre-exposed to the Epac inhibitor ESI-09 (10 µM, Fig. 2B and C), confirming that adiponectin exocytosis is stimulated via activation of this intracellular cAMP receptor (Komai *et al*., 2016). Consistent with previous studies (El Hachmane *et al*., 2015; Komai *et al*., 2014; Komai *et al*., 2016), exocytosis was stimulated by a pipette solution containing 0.1 mM cAMP and a low concentration of the Ca^2+^ chelator EGTA (50 μM; IC-2) and the peak rate of exocytosis (measured during the second minute) averaged 7.0±1.6 fF/s. Addition of ATP to adipocytes infused with IC-2 increased ΔC/Δt by ∼40% (Fig. 2D and E). Collectively, our results demonstrate that NA triggers adiponectin exocytosis via the Epac signalling pathway and that ATP potentiates cAMP-triggered adiponectin exocytosis. In addition, ATP can stimulate adiponectin secretion in the absence of a concomitant cAMP elevation.

### Ca^2+^-dependent mechanisms are involved in ATP-stimulated but not in NA-triggered adiponectin secretion

We next investigated the involvement of Ca^2+^ in the adiponectin secretion stimulated by ATP or NA. Extracellularly applied ATP elevates white adipocyte [Ca^2+^]_i_ (Kelly *et al*., 1989; Lee *et al*., 2005) and our own findings have demonstrated that ATP increases 3T3-L1 adipocyte [Ca^2+^]_i_ via activation of purinergic P2Y2 receptors (P2Y2Rs) and ensuing store-operated Ca^2+^ entry (SOCE; El Hachmane *et al*., 2018; El Hachmane and Olofsson, 2018). As shown in Fig. 3A, ATP stimulated adiponectin secretion >2-fold over basal in 3T3-L1 adipocytes (30 min static incubation). Adiponectin release tended to still be elevated by ATP in Ca^2+^-depleted cells (pre-treatment with the Ca^2+^ chelator BAPTA; P=0.1). To more specifically determine effects of ATP on adiponectin vesicle replenishment (Komai *et al*., 2014), BAPTA-treated and untreated 3T3-L1 adipocytes were incubated with ATP during 60 minutes. ATP alone stimulated adiponectin release 1.6-fold in this series of experiments. The secretion was not significantly elevated over basal (control) in Ca^2+^-chelated cells (Fig. 3B). BAPTA slightly inhibited adiponectin secretion stimulated by a combination of the two secretagogues (Fig. 3C), thus in line with an effect of Ca^2+^ buffering on the ATP but not NA signalling pathway. Ratiometric Ca^2+^ imaging in Fura-2 loaded 3T3-L1 adipocytes confirmed the ability of ATP to elevate [Ca^2+^]_i_ (El Hachmane *et al*., 2018; Kelly *et al*., 1989; Lee *et al*., 2005), from a basal value of 146±3 nM to a peak averaging 307±7 nM (P<0.01) whereas ATP was without effect on [Ca^2+^]_i_ in BAPTA-treated cells (Fig. 3D). NA did not affect [Ca^2+^]_i_ in 3T3-L1 adipocytes and the [Ca^2+^]_i_ averaged 174±4 nM before and 177±4 nM 2 min after addition of the catecholamine; the responsiveness of the cells was confirmed by addition of ATP at the end of the recordings (Fig. 3E).

**Fig. 3.**
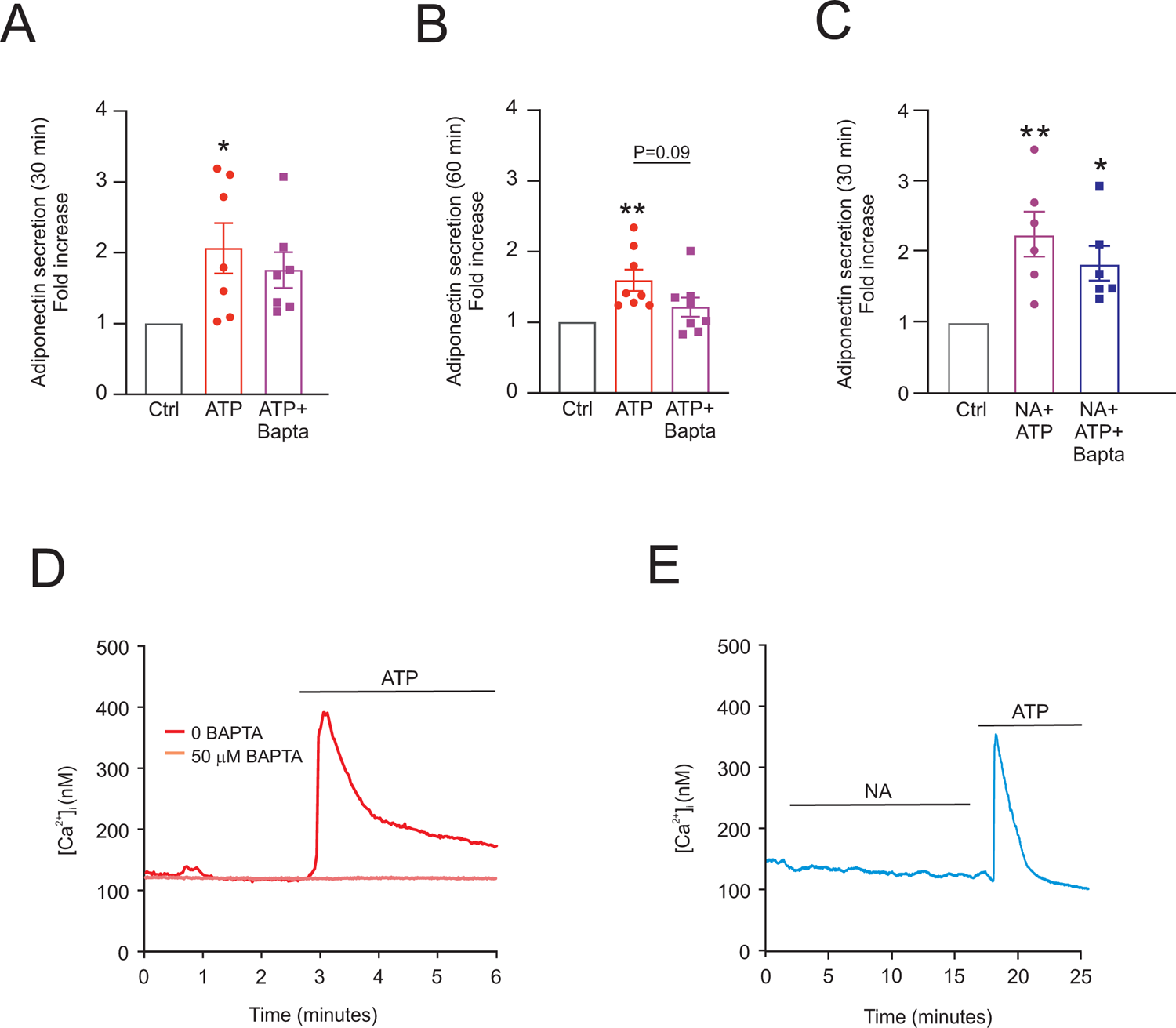
The role of cytosolic calcium in NA- and ATP-stimulated adiponectin secretion. Adiponectin release in 3T3-L1 adipocytes during 30 min (***A***) and 60 min (***B***) incubations with ATP (100 µM) with or without BAPTA pre-treatment (50 µM for 30 min). ***C***, Adiponectin release in 3T3-L1 adipocytes during 30 min incubations with NA (100 nM) in combination with ATP (100 µM), with or without BAPTA pre-treatment (50 µM for 30 min). ***D***, Example traces of the effect of extracellular ATP on [Ca^2+^]_i_ in 3T3-L1 adipocytes with or without BAPTA pre-treatment (50 µM for 30 min). Results in ***A*** - ***C*** are from 7-8 experiments and expressed as fold-increase compared to control (5 mM glucose). Recordings in ***D*** are representative of 5 separate experiments in the absence of BAPTA and 162 individually analysed cells. Analysis of BAPTA-treated cells was performed on averaged curves from 4 separate experiments from a total of 101 cells; the [Ca^2+^]_i_ averaged 130±13 nM before and 134±10 nM after ATP-application. ***E***, Example trace of the effect of NA (100 nM) on [Ca^2+^]_i_ in 3T3-L1 adipocytes, representative of 5 separate experiments and 89 individually analysed cells. *\*P<0.05; **P<0.01*.

To in more detail investigate the role of Ca^2+^ in ATP-stimulated exocytosis, we performed capacitance recordings with one Ca^2+^-depleted pipette solution (buffered with 10 mM EGTA) and one pipette solution that contained a high free concentration of Ca^2+^ (∼1.5 μM). Both solutions were supplemented with cAMP (IC-3 and IC-4, respectively). Thus, IC-3 rapidly buffers a Ca^2+^ increase and IC-4 clamps the concentration of Ca^2+^ at a high enough level to mask endogenous elevations of [Ca^2+^]_i_. ATP was without effect on exocytosis triggered by either solution (Fig. 4A and B). In an additional series of capacitance recordings, ATP was added to cells infused with IC-1 (the cAMP- and Ca^2+^-depleted solution containing 10 mM BAPTA). The response to external ATP was heterogeneous under those conditions; the nucleotide clearly triggered a small magnitude of exocytosis in >60% of investigated adipocytes (Fig. 4C) while remaining cells were unaffected (Fig. 4D). ADR was added at the end of each experiment to verify that cells were responsive (Komai *et al*., 2016). Analysis of average rates including all experiments showed a small but significant ATP stimulation of exocytosis and ΔC/Δt averaged 0.01±0.4 fF/s before application of ATP *vs.* 2.7±0.8 fF/s (P<0.01) and 2.9±0.8 fF/s (P<0.01) 1 and 2 minutes after application of the nucleotide respectively.

**Fig. 4.**
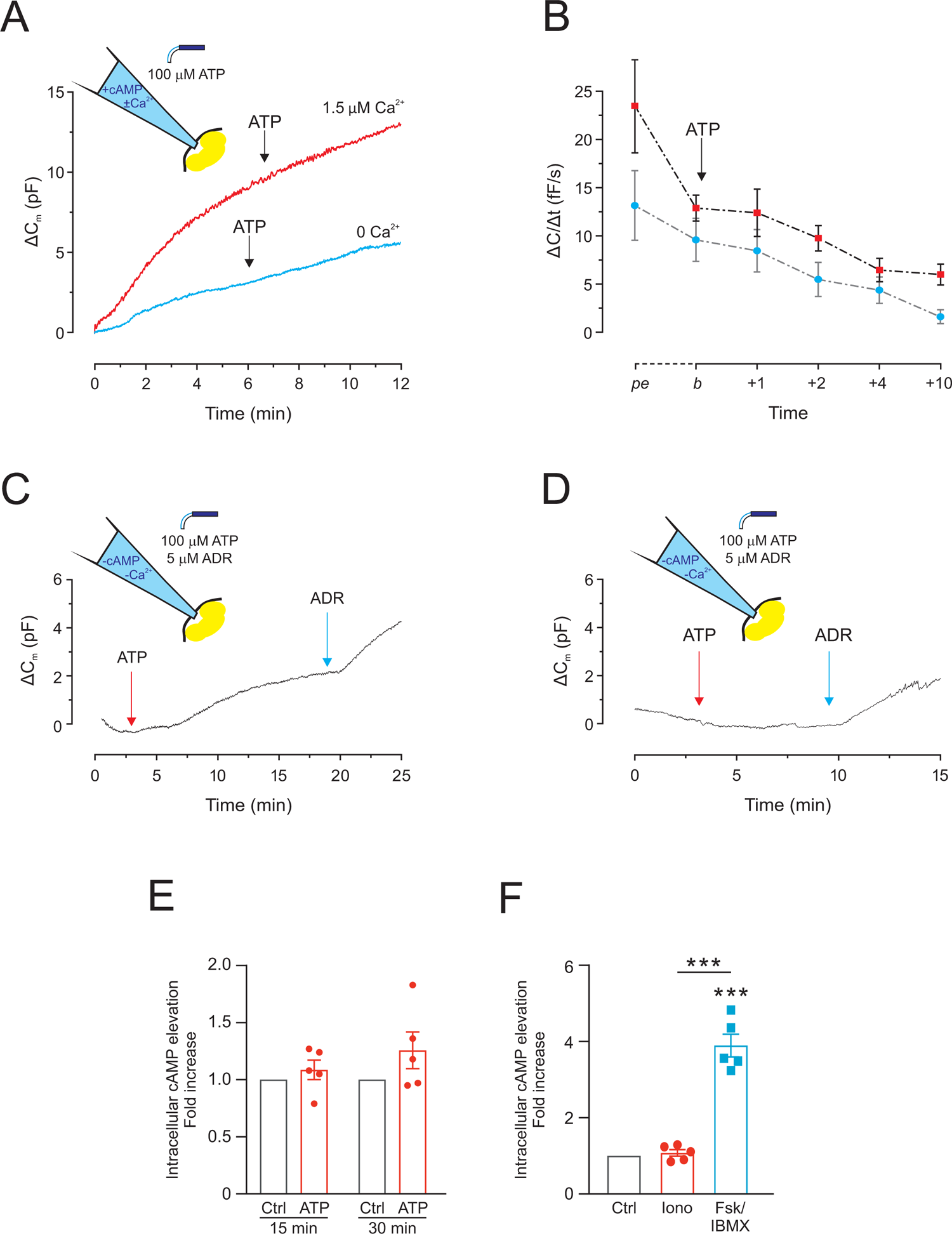
The role of cAMP in ATP-stimulated adiponectin secretion. ***A,*** Two representative traces of Δ*C*_m_ for cells dialyzed with either a Ca^2+^-depleted pipette solution containing cAMP (IC-3) or a solution with a combination of cAMP and 1.5 µM free Ca^2+^ (IC-4). ***B,*** Average Δ*C*/Δ*t* is given at indicated time points. Time point “*pe*” is the *pe*ak rate measured during the second minute after start of the recording (thus during peak exocytosis). *“b”* indicates the time point just *b*efore application of ATP and additional time-points are measured between 1 and 10 minutes after *b* (+1 – +10). ***C*** and ***D,*** Example traces showing the heterogeneous effect of external ATP, followed by addition of ADR (to ascertain cell responsiveness), as indicated. Cells were infused with a non-stimulatory solution lacking cAMP together with 10 mM BAPTA (IC-1). ***E***, Intracellular cAMP levels in 3T3-L1 adipocytes in response to ATP-treatment (15 or 30 min). ***F***, Levels of cAMP in 3T3-L1 adipocytes treated for 30 min with Ionomycin (5µM) or a combination of Forskolin (10 nM) and IBMX (200 µM). Data in ***B*** are from 5 (1.5 µM Ca^2+^) and 7 (0 Ca^2+^) recordings. Traces in ***C*** and ***D*** are representative of 6 and 3 recordings respectively. Results in ***E*** and ***F*** are expressed as fold-increase compared to control (5 mM glucose) and represent 5 experiments *\*\*\*P<0.001*.

Collectively, the secretion and capacitance data suggest that the ATP-induced adiponectin exocytosis chiefly depend on an elevation of [Ca^2+^]_i_, but that ATP also may exert Ca^2+^-independent effects. An elevation of [Ca^2+^]_i_ can lead to increased production of cAMP via activation of Ca^2+^-sensitive adenylyl cyclases (Halls and Cooper, 2011). Moreover, extracellular ATP has been shown to also induce production of cAMP via Ca^2+^-independent pathways (Anwar et al., 1999; van der Weyden et al., 2000). To test the hypothesis that ATP affects adipocyte cAMP levels, we measured intracellular cAMP in cells exposed to ATP during 15 or 30 minutes. As shown in Fig. 4E, the cAMP levels were similar in adipocytes incubated in the presence or absence of ATP (although ATP tended to increase cAMP in 30 min incubations; P=0.15). To more directly determine the effects of an [Ca^2+^]_i_ increase on intracellular cAMP, 3T3-L1 adipocytes were incubated with the Ca^2+^ ionophore ionomycin that potently elevates adipocyte [Ca^2+^]_i_ (El Hachmane *et al*., 2015). Ionomycin (1 μM) stimulated adiponectin secretion, although not as effectively as FSK/BIX (a 1.3±0.2 fold increase over basal with ionomycin compared with 1.9±0.5 fold for FSK/BIX; P<0.01 for both *vs.* control; n=5; not shown). FSK/IBMX elevated cAMP levels ∼4-fold whereas cellular cAMP levels were, in agreement with published data (Musovic et al., 2021), unaffected by ionomycin (Fig. 4F).

Together, our data suggest that the ability of ATP to induce adiponectin exocytosis is largely due to an elevation of [Ca^2+^]_i_ while NA effects are Ca^2+^-independent. In addition, ATP stimulates a smaller magnitude of adiponectin exocytosis/secretion via Ca^2+^-independent mechanisms.

### The ATP-stimulated adiponectin secretion is blunted in adipocytes isolated from mice with diet-induced obesity

We have previously demonstrated that adrenergically triggered adiponectin secretion is diminished in adipocytes isolated from mice with diet-induced obesity/diabetes (Komai *et al*., 2014). To determine if ATP-stimulated adiponectin release is likewise disturbed in metabolic disease, we incubated primary IWAT adipocytes isolated from mice fed chow-(“lean adipocytes”) or HFD diet (“obese adipocytes”) with ATP in the presence or absence of NA. The HFD-fed mice were obese (total body weight averaged 29.5±1.2 g and 42.6±0.6 g for chow- and HFD-fed animals respectively; P<0.001) and diabetic as demonstrated by elevated serum insulin (Fig. 5A) and blood glucose (Fig. 5B) levels. The IWAT weight was increased in obese compared to lean mice (Fig. 5C). In agreement with results in Fig. 2A, NA or ATP alone stimulated adiponectin secretion in lean adipocytes and the combination of the catecholamine and the nucleotide triggered adiponectin release significantly more potent than ATP alone (P=0.1 *vs.* NA only). However, NA or ATP alone or in combination were unable to stimulate adiponectin release in obese adipocytes (Fig. 5D). The cellular adiponectin levels were similar in adipocytes isolated from lean and obese mice and averaged 1.4±0.1 and 1.6±0.2 mg/g protein respectively (P=0.4; n=15) and the percentage of released adiponectin (out of total content) was significantly lower in the HFD group (Fig. 5E). The results in Fig. 5D and E jointly propose that the blunted adiponectin release is due to a secretion defect and not a result of diminished adiponectin synthesis.

**Fig. 5.**
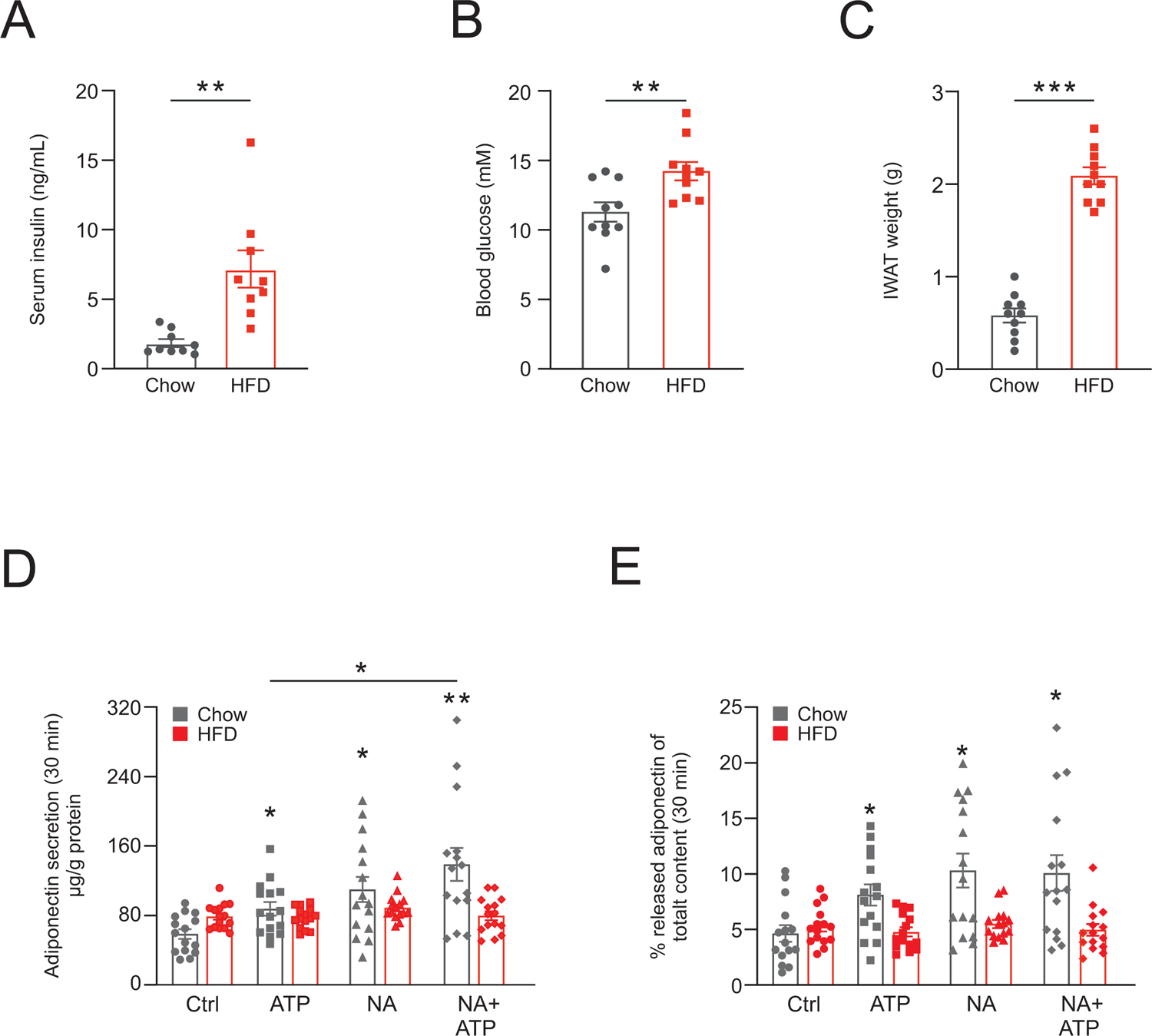
Purinergic and adrenergic stimulation of adiponectin secretion using adipocytes isolated from lean and obese/diabetic mice. Serum insulin (***A***) and blood glucose (***B***) levels in mice fed chow or HFD through 8 weeks. ***C,*** Subcutaneous adipose tissue (IWAT) mass obtained from chow- and HFD-fed mice. ***D,*** Adiponectin secretion (30 min) in primary IWAT adipocytes isolated from chow- or HFD-fed mice. Cells were incubated together with ATP, NA or a combination of NA and ATP as indicated. ***E,*** Released adiponectin expressed as percentage of total adipocyte adiponectin content. Results in ***A-C*** are from 9-10 chow or HFD mice. Results in ***D*** and ***E*** are from 15 experiments with adipocytes isolated from 5 chow or HFD fed mice. *\*P<0.05*; *\*\*P<0.01*; *\*\*\*P<0.001*.

Our data demonstrates that the purinergic signalling is disturbed in obese adipocytes and confirm the previously reported presence of catecholamine resistance in obese/diabetic mice (Komai *et al*., 2016).

### Molecular characterisation of the ATP signalling pathway involved in the control of adiponectin secretion

We have previously determined the molecular details of the adrenergic signalling pathway involved in catecholamine-triggered adiponectin exocytosis. In this work we defined the necessity of β_3_ARs and Epac1 for adrenaline-stimulated adiponectin secretion by individual siRNA knockdown of the two proteins. We also demonstrated that β_3_ARs and Epac1 protein levels are decreased by ∼30% in obese IWAT adipocytes (Komai *et al*., 2016). Therefore, we next characterised the signalling pathways involved in the purinergic regulation of adiponectin secretion in more detail. Consistent with previous studies (Laplante *et al*., 2010; Lee *et al*., 2005; Omatsu-Kanbe et al., 2006), P2Y2Rs were readily expressed in mouse IWAT adipocytes. Interestingly, the expression of P2Y2Rs was markedly reduced in adipocytes isolated from obese animals, both at the gene (Fig. 6A) and protein (Fig. 6B) level. In agreement with published data (Komai *et al*., 2016), gene expression of β_3_ARs and Epac1 were lower in obese compared to lean adipocytes (a decrease by >50% for β_3_AR and >70% for Epac1; Supplementary Fig. 1). ATP has been show to elevate 3T3-L1 adipocyte [Ca^2+^]_i_ via activation of P2Y2Rs (El Hachmane *et al*., 2018; El Hachmane and Olofsson, 2018) but ATP effects on adiponectin release were not investigated in this work. We treated lean IWAT adipocytes with the P2Y2R antagonist AR-C 118925XX (AR-C; 10 µM). NA combined with ATP stimulated adiponectin secretion >2-fold in un-treated adipocytes; the release was still triggered in the presence of AR-C, but of a magnitude significantly decreased compared to control cells (Fig. 6C). P2Y2Rs couple to the G protein Gq11 and elevates [Ca^2+^]_i_ via phospholipase C (PLC) to generate the second messengers inositol 1,4,5-triphosphate (IP3) and diacylglycerol (DAG; Taylor and Tovey, 2010). Pre-treatment of adipocytes with the PLC inhibitor U73122 abrogated the ATP-induced [Ca^2+^]_i_ increase. In cells treated with U73122, the [Ca^2+^]_i_ averaged 139±4 nM 2 min before and 139±3 nM after ATP-application whereas the [Ca^2+^]_i_ averaged 101±3 nM before and 340±9 nM 2 min prior and following the addition of ATP in cells treated with the inactive analogue U73343. ATP still stimulated adiponectin release in U73343-treated adipocytes while secretion was blunted in cells pre-exposed to U73122 (Fig. 6E).

**Fig. 6.**
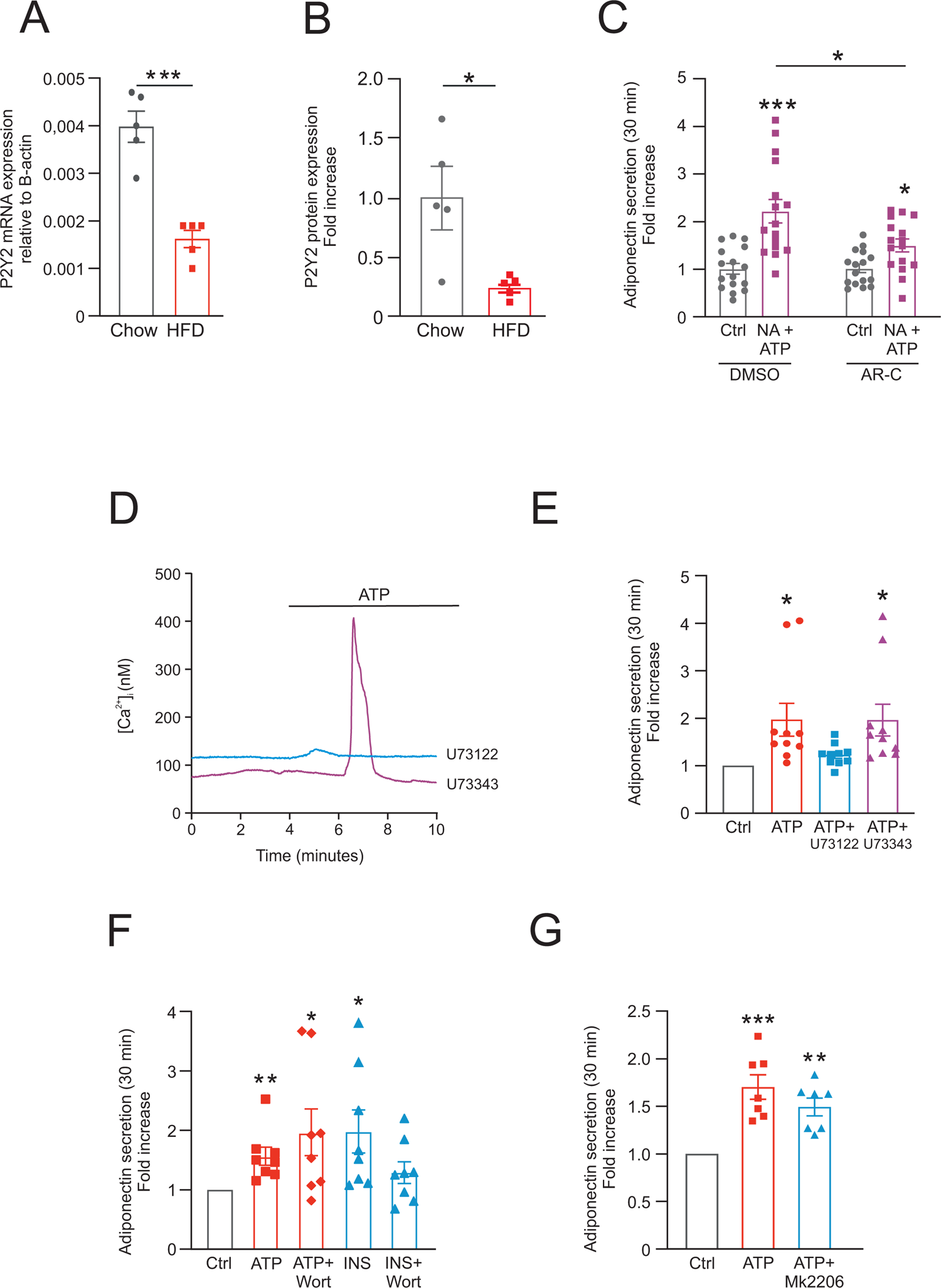
Signalling pathways involved in ATP-stimulated adiponectin secretion. Gene (***A***) and protein expression analysis of P2Y2 receptor in IWAT adipocytes isolated from chow- and HFD-fed mice. Gene expression of the P2Y1 receptor was also detected at a low level, but the expression was unaltered in adipocytes from obese mice (not shown). ***C,*** Effects of pre-treatment with P2Y2 receptor antagonist AR-C (1 µM) for 30 min on the NA/ATP-stimulated adiponectin release in IWAT adipocytes. ***D***, Example traces of the effect of extracellular ATP on [Ca^2+^]_i_ in 3T3-L1 adipocytes in the presence of the PLC inhibitor U73122 (10 μM) or the inactive analogue U73343 (10 μM). ***E***, ATP-stimulated adiponectin release (30 min) in 3T3-L1 adipocytes pre-treated with U73122 or U73343. ***F,*** ATP-stimulated adiponectin secretion (30 min) in control (Ctrl) 3T3-L1 adipocytes or 3T3-L1 adipocytes pre-treated with PI3K-inihibitor Wortmannin (100 nM during 30 min). ***G,*** ATP-stimulated adiponectin secretion (30 min) in control (Ctrl) or 3T3-L1 adipocytes pre-treated with the PKB/AKT2-inihibitor Mk2206 (1 µM during 30 minutes). Data in ***A*** and ***B*** are from 5 chow and 5 HFD mice. Results in ***C*** are from 16 experiments in IWAT adipocytes isolated from 7 mice. Recordings in ***D*** are representative of 6 separate and 64 individually analysed cells. Results in ***E, F*** and ***G*** are expressed as fold-increase compared to control (5 mM glucose) and represent 7-10 experiments. Gene expression was normalised against β-actin (*Actb*) using the relative 2^ΔC_t_ method. Primers were used at a concentration of 500 nM. *\*P<0.05*; *\*\*P<0.01*; *\*\*\*P<0.001*.

ATP signalling via P2Y2Rs can also activate the phosphoinositide 3-kinase (PI3K) Akt signalling pathway (Katz et al., 2011). Since adiponectin release can be induced by insulin (Brannmark *et al*., 2020), via PI3K-dependent mechanisms (Cong et al., 2007; Lim et al., 2015), we hypothesized that PI3K signalling might partake in ATP-stimulated adiponectin secretion. However, ATP remained capable of stimulating adiponectin release in 3T3-L1 adipocytes pre-treated with the PI3K inhibitor Wortmannin. If anything, secretion tended to be potentiated in adipocytes treated with the inhibitor. It was confirmed that Wortmannin inhibited insulin-induced adiponectin release (Fig. 6F). ATP likewise stimulated secretion of adiponectin in cells treated with the Akt inhibitor MK-2206 (Fig. 6G).

Collectively, our data proposes that ATP stimulates adiponectin secretion via the P2Y2/Gq11/PLC pathway that elevates [Ca^2+^]_i_, with no involvement of PI3K/Akt signalling.

## Discussion

Here we show that sympathetic innervation is disturbed in adipose tissue from mice with diet-induced obesity, demonstrated by largely reduced levels of adipose tissue tyrosine hydroxylase (TH) and noradrenaline (NA), and that this is associated with decreased serum HMW/total adiponectin. With the aim to determine the role of sympathetic innervation for the pathophysiological regulation of adiponectin secretion at a molecular level, we have investigated how NA and ATP, alone or together, affect adiponectin exocytosis/secretion in cultured 3T3-L1 and in primary subcutaneous (IWAT) mouse adipocytes. Our results demonstrate that NA potently stimulates adiponectin exocytosis via cAMP and Epac1 and that ATP chiefly induces adiponectin release via activation of P2Y2Rs and an elevation of [Ca^2+^]_i_. An important role of ATP appears to be to potentiate the NA-triggered adiponectin secretion, although ATP in addition stimulates release of the adipocyte hormone in the absence of a concomitant cAMP increase. Based on our findings, we propose a central role for SNS innervation in the control of adiponectin release and that an interplay between adrenergic and purinergic signalling partakes in the regulation of adiponectin secretion. Some key results and their relation to adipocyte function and whole body pathophysiology are conferred below.

### Noradrenaline and ATP jointly control adiponectin secretion

White adipocyte adiponectin exocytosis is stimulated by β_3_AR-mediated increases of intracellular cAMP and can be augmented by a simultaneous elevation of [Ca^2+^]_i_ (El Hachmane *et al*., 2015; Komai *et al*., 2014; Komai *et al*., 2016). The profuse sympathetic innervation of white adipose tissue (Bartness *et al*., 2010) together with the knowledge that β_3_ARs have a >30-fold higher affinity for NA compared to adrenaline (ADR; Lohse et al., 2003), suggest NA as a chief regulator of adiponectin secretion. Moreover, the NA levels in rat IWAT are, as can be expected for a locally distributed compound, 80-fold higher than ADR (secreted by the adrenal medulla; Vargovic et al., 2011). Indeed, NA triggered a peak rate of adiponectin exocytosis comparable to that attained upon stimulation with ADR (*c.f.* Fig. 2C in the current work with Fig. 2D of Komai *et al*., 2016) and the two catecholamines stimulated secretion of similar amounts of adiponectin (*c.f.* Fig. 2A in the present study with Fig. 1A of Komai *et al*., 2016). The signalling pathway responsible for the Ca^2+^ potentiation of catecholamine/cAMP-stimulated adiponectin release (El Hachmane *et al*., 2015; Komai *et al*., 2014) has remained unknown. Catecholamines can themselves elevate adipocyte [Ca^2+^]_i_ via activation of α1 adrenergic receptors (Komai *et al*., 2016; Seydoux et al., 1996). We have shown that the slight [Ca^2+^]_i_ increase induced by ADR is without effect on adiponectin secretion (Komai *et al*., 2016) and the data in Fig. 3E demonstrates that the adipocyte [Ca^2+^]_i_ was unaffected by NA. Thus, the signalling underlying the Ca^2+^-augmentation of cAMP-stimulated adiponectin exocytosis (El Hachmane *et al*., 2015; Komai *et al*., 2014) must involve other molecular components. ATP increases [Ca^2+^]_i_ in white adipocytes (El Hachmane *et al*., 2018; Kelly *et al*., 1989; Laplante *et al*., 2010; Lee *et al*., 2005). The fact that ATP is co-released with NA from sympathetic nerves (Bartness *et al*., 2010) lead us to hypothesise that purinergic signalling is involved in the Ca^2+^-dependent augmentation of adiponectin secretion. Indeed, the results of the current study support that sympathetic innervation controls white adipocyte adiponectin exocytosis in agreement with the model presented in Fig. 7: As depicted in the left part of Fig. 7, NA, released from sympathetic nerves, binds to β_3_ARs which leads to an elevation of cAMP and activation of Epac1 (Komai *et al*., 2016) whereupon release-ready adiponectin-containing vesicles fuse with the adipocyte plasma membrane, to discharge their cargo to the extracellular space. The co-secreted ATP binds to P2Y2Rs in the adipocyte plasma membrane, to increases [Ca^2+^]_i_ that will induce further adiponectin exocytosis. The secretion model proposes that external ATP is important for the previously demonstrated Ca^2+^-dependent maintenance of adiponectin release over longer time-periods, corresponding to replenishment of release-ready adiponectin vesicles (Komai *et al*., 2014). However, our data show that Ca^2+^ buffering affects both early (fusion of release-ready vesicles in Fig. 3A) and prolonged (Fig. 3B) secretion of adiponectin, although the effect on early release appears to be smaller. This dual effect of ATP-induced [Ca^2+^]_i_ is in agreement with previous work, demonstrating that Ca^2+^, in addition to its role in vesicle replenishment, also augments the rapid cAMP-triggered exocytosis (Komai *et al*., 2014).

**Fig. 7.**
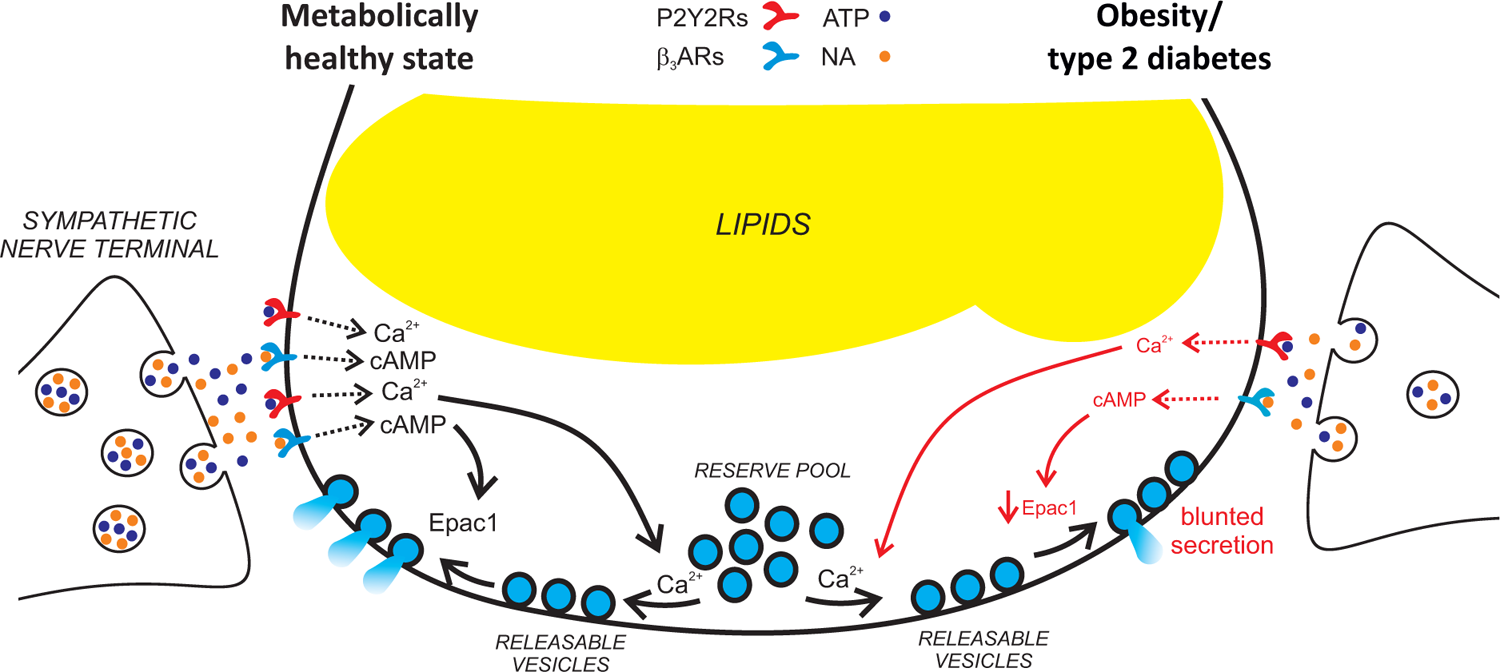
Model of proposed physiological regulation of adiponectin exocytosis in metabolic health and disease. ***Left part:*** Exocytosis of adiponectin containing vesicles is stimulated by NA-binding to β_3_ARs and downstream activation of Epac1. Activation of P2Y2 receptors by co-secreted ATP potentiates the adrenergically stimulated adiponectin release through elevations of [Ca^2+^]_i._. ***Right part:*** The adrenergic stimulation as well as the purinergic augmentation of adiponectin exocytosis is blunted in obesity/type 2 diabetes due to lower levels of β_3_ARs and Epac1 as well as reduced expression of P2Y2 receptors. *See text for more details*.

### Decreased adrenergic and purinergic signalling in obesity-associated diabetes leads to blunted adiponectin secretion and reduced serum HMW adiponectin

Our current study confirms earlier findings of that catecholamine-stimulated adiponectin release is attenuated in the obese/diabetic state, due to reduced abundance of β_3_ARs in IWAT adipocytes (Komai *et al*., 2016). However, we now show that also the purinergic signalling pathway is disrupted in adipocytes isolated from mice with diet-induced obesity, as demonstrated by a ∼70% reduction of P2Y2R protein levels (Fig. 6B), connected with blunted ATP-induced adiponectin release (Fig. 5D and E). We propose that obesity-associated diabetes leads to abrogated exocytosis of adiponectin-containing vesicles as portrayed in the right part of Fig. 7. In our model, the lower abundance of β_3_ARs dampens the cAMP increase, as previously demonstrated (Komai *et al*., 2016). We now add to our secretion model that the lower abundance of P2Y2Rs results in reduced Ca^2+^ potentiation of adiponectin exocytosis. The lower levels of NA in obese white adipose tissue (Fig. 1C) likely further aggravates the pathological state. We expect that the concentration of ATP is likewise lower in the interstitial compartment within adipose tissue; this is unfortunately not possible to measure since ATP is present at high concentrations within cells. The proposed importance of adrenergic and purinergic signalling for the pathophysiological control of adiponectin secretion is supported by the observed reduction of circulating HMW adiponectin in mice with obesity-associated diabetes (Fig. 1D and Komai *et al*., 2016). Of interest, volume fluorescence-imaging studies have revealed a dense network of sympathetic fibres in close proximity to more than 90% of adipocytes within white adipose tissue (Jiang et al., 2017), in support of that sympathetic innervation of individual adipocytes is essential for the regulated adiponectin exocytosis.

### Adiponectin secretion stimulated by ATP alone

The finding that ATP alone stimulates adiponectin secretion/exocytosis when cAMP is not concomitantly elevated may appear puzzling. Our own published work shows that Ca^2+^ is unable to induce exocytosis in adipocytes in the absence of cAMP (depleted from the pipette solution during patch-clamping) and that the cAMP-triggered secretion of adiponectin is unaffected by Ca^2+^-depletion (BAPTA buffering; Komai *et al*., 2014; Komai *et al*., 2016). However, it is important to recall that whereas the intracellular milieu of cells used for electrophysiological studies is clamped by the constituents in the pipette solution, the adipocytes used for secretion data are metabolically intact and therefore able to produce endogenous cAMP. The basal (non-stimulating) levels of cAMP can likely act together with the nucleotide-induced [Ca^2+^]_i_ elevation to prompt adiponectin secretion. This proposal is supported by that the Ca^2+^ ionophore ionomycin on its own stimulates adiponectin release in intact 3T3-L1 adipocytes, without affecting cAMP-levels (Fig. 4F).

ATP was found to also stimulate adiponectin exocytosis via Ca^2+^-independent effects (Fig. 4C). Extracellularly applied ATP has been shown to prompt synthesis of cAMP (Anwar *et al*., 1999; Halls and Cooper, 2011; van der Weyden *et al*., 2000), but we were unable to detect significantly elevated cAMP levels in adipocytes exposed to ATP (Fig. 4E) or ionomycin (Fig. 4F). It should be noted though that GPCRs are typically organised in micro-domains. This spatial organisation facilitates communication between different receptor types and aids interaction between receptors and their intracellular target signalling proteins, making cAMP and Ca^2+^ signalling constrained to specific regions (Willoughby, 2012; Willoughby and Cooper, 2007). Thus, it is possible that ATP increases cAMP in confined sub-plasma membrane exocytotic regions (supported by the tendency to an elevation of cAMP in cells exposed to ATP through 30 min; Fig. 4E) but that this can not be measured with the applied technique (where total cAMP is analysed in the cell homogenate). An alternative explanation for the Ca^2+^-independent effect of ATP on adiponectin release is activation of small GTPases, a type of G-proteins that are important regulators of exocytosis (Wu et al., 2008). P2Y2R signalling activates Rac and Rho GTPases (Erb et al., 2006). Importantly though, our combined data proposes that the ATP-induced adiponectin exocytosis predominantly depend on Ca^2+^; ATP potentiates cAMP-stimulated exocytosis when cytoplasmic Ca^2+^ is allowed to fluctuate (Fig. 2D and E) but neither when Ca^2+^ is kept high, nor when Ca^2+^ is strongly buffered (Fig. 4A and B).

### ATP in adipose tissue

Extracellular ATP has been shown to regulate both glucose uptake (Adamson et al., 2015) and lipolysis (Tozzi et al., 2019) in adipose tissue. In addition to its release from sympathetic nerve terminals (Bartness *et al*., 2010), ATP is co-secreted via vesicular release from a number of endocrine cell types (Detimary et al., 1996; Leitner et al., 1975; Rojas et al., 1985; Winkler and Westhead, 1980). Also non-excitable cell types such as thrombocytes and epithelial cells release substantial amounts of the nucleotide (reviewed in Praetorius and Leipziger, 2009). ATP is likely released via vesicular exocytosis also in white adipocyte, although this has never been shown. It is of interest that recent studies demonstrate that murine adipocytes release ATP via a channel-mediated mechanism involving pannexin-1 (Adamson *et al*., 2015; Tozzi *et al*., 2019). The ATP-permeable pannexin-1 pore has been demonstrated to be regulated via cAMP/PKA-dependent mechanisms (Tozzi *et al*., 2019), thus emphasising the adrenergic-purinergic signalling crosstalk. Incorporated into our adiponectin exocytosis model, the catecholamine-stimulation will not only trigger secretion of adiponectin-containing vesicles, but the cAMP elevation will in addition stimulate release of ATP via pannexin-1. The secreted ATP can consequently augment adiponectin release in an autocrine and paracrine fashion. Regardless of the source (sympathetic nerves or the adipocyte itself), ATP is released in close/direct vicinity to the adipocyte plasma membrane and can thus act at locally high concentrations. The fact that ATP secretion may be confined to restricted regions (Allen et al., 2007; Erb and Weisman, 2012; Insel et al., 2005) make it difficult to reliably measure the local concentration acting on purinergic receptors. However, quantification of near-membrane peak concentrations of ATP by use of biosensors has demonstrated local concentrations in the ten to hundred μM range (Praetorius and Leipziger, 2009). ATP is rapidly hydrolysed and produced adenosine can bind to A1-adenosine receptor and instead decrease intracellular cAMP levels. Adenosine has been shown to exert an antilipolytic effect (Fredholm, 1981) and this proposes that ATP, or rather its break-down products, might also functions as a negative feedback regulator, of adiponectin exocytosis and of other metabolic processes in the white adipocytes.

### The sympathetic nervous system and metabolic disease – future perspectives

The regulation of energy homeostasis is controlled by the SNS and both sympathetic and parasympathetic branches have been proposed to contribute to the development of obesity and metabolic dysfunction (Burnstock and Gentile, 2018). The obesity-associated changes of SNS activity are not fully clarified and different studies have reported reduced, increased or unaltered SNS responsiveness. Moreover, the obesity-associated alterations of SNS innervation or activity are not uniform in different tissues and organs. Nonetheless, several studies appear to agree on that SNS innervation/activity of adipose tissue is upregulated in obese subjects under resting conditions while the responsiveness to sympathetic stimulation is dampened (Bartness *et al*., 2014; Davy and Orr, 2009; Tentolouris et al., 2006). In support of this, obesity has been linked to increased basal lipolysis in adipose tissue, while catecholamine-stimulated lipolysis is reduced (Duncan et al., 2007). Enhanced sympathetic activity in the resting state, resulting in elevated basal NA and ATP levels within white adipose tissue, can be envisaged to render adipocytes unresponsive to both adrenergic and purinergic signalling. GPCRs are known to be desensitized upon over-activation and prolonged overstimulation leads to receptor sequestration, degradation and reduced gene expression (Black et al., 2016; Rajagopal and Shenoy, 2018). Adipocyte function may thus be compromised in a similar way to what we observe here. Over time, the obesity-associated disturbance of sympathetic innervation might exacerbate the pathophysiological situation and further reduce the stimulated adiponectin secretion. It would be of great interest to study the changes of adiponectin secretion and circulating levels during development of obesity and metabolic disease as well as possible parallel alterations of SNS activity. It is intriguing to consider that early interventions aimed at reducing obesity/diabetes-associated increases of basal SNS activity may result in largely sustained adipocyte responsiveness to adrenergic and purinergic signalling and deter the hypoadiponectinemia observed in obesity-associated metabolic disease. In light of the global obesity epidemic, the need to develop novel anti-obesity drugs is evident. Therapeutic purinergic signalling approaches have been proposed for the treatment of obesity and its related comorbidities (Burnstock and Gentile, 2018). The observation that extracellular ATP controls both glucose uptake (Adamson *et al*., 2015) and lipolysis (Tozzi *et al*., 2019) together with the findings presented here, suggest that targeting purinergic signalling pathways in adipocytes might be a feasible future approach to manage obesity and metabolic dysfunction.

## Additional information

## Data availability

All data generated or analysed during this study are included in this published article or as supplementary files.

## Author contributions

Conception and design of the experiments: S.M., A.M.K. and C.S.O. Data collection, analysis and interpretation of data S.M., A.M.K., M. K. S., Y.W., I.W.A. and C.S.O. Writing of the manuscript C.S.O. and S.M. All authors drafted and revised the manuscript as well as read and approved of the final version. All experiments were carried out at the Department of Physiology/Metabolic Physiology, Gothenburg University. C.S.O. is the guarantor of this work and, as such, had full access to all the data in the study and takes responsibility for the integrity of the data and the accuracy of the data analysis.

## Acknowledgements

We are grateful for the help provided by Peter Micallef, Seid Talavanic and Ali Abdali (Department of Physiology/Metabolic Physiology).

---

**Supplementary Fig. 1.**
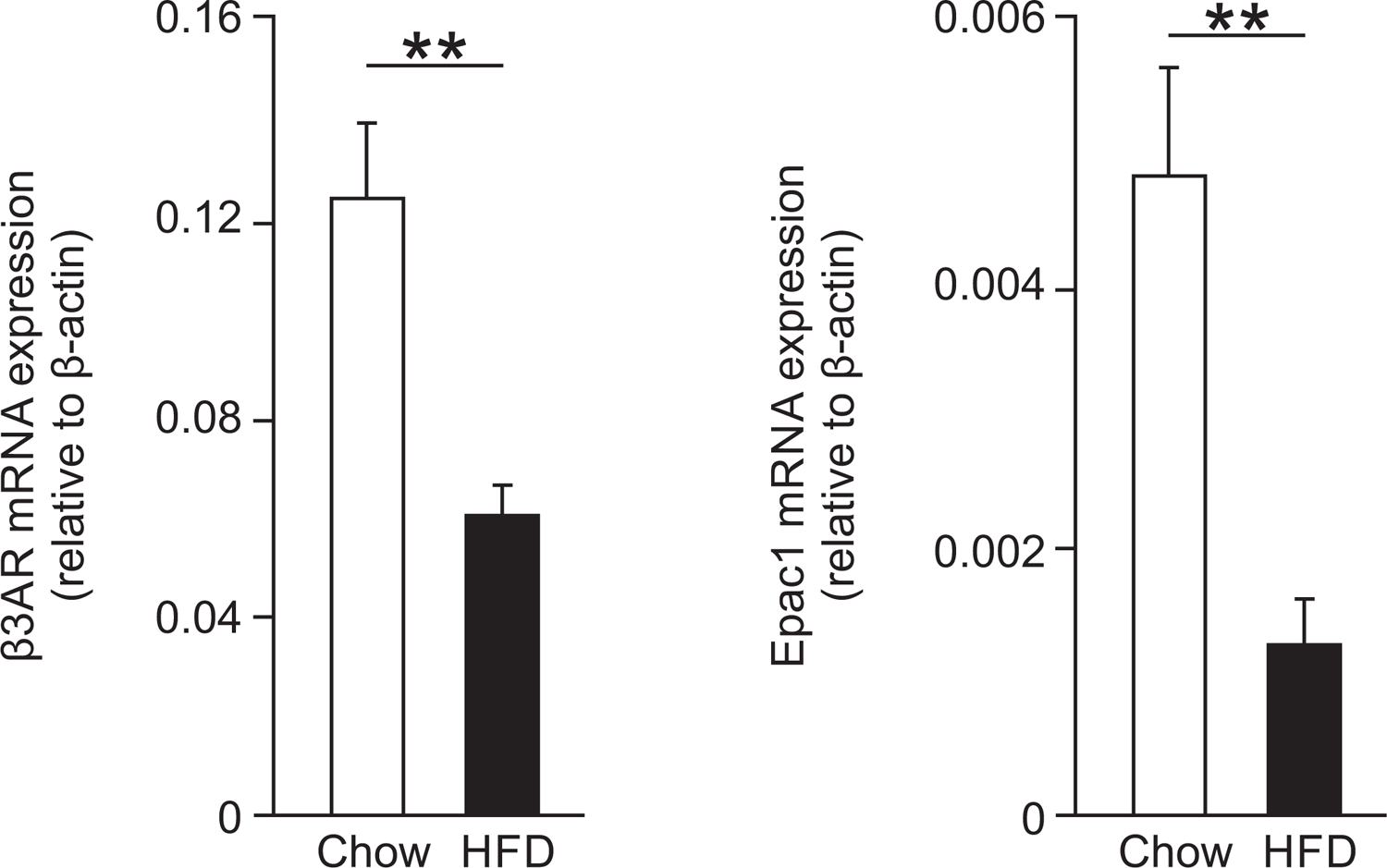
Decreased gene expression of β3ARs and Epac1in adipocytes isolated from obese (HFD-fed) mice, compared to adipocytes from lean (chow-fed) animals. Results from 5 chow and 5 HFD mice.

**Supplementary Table 1.**
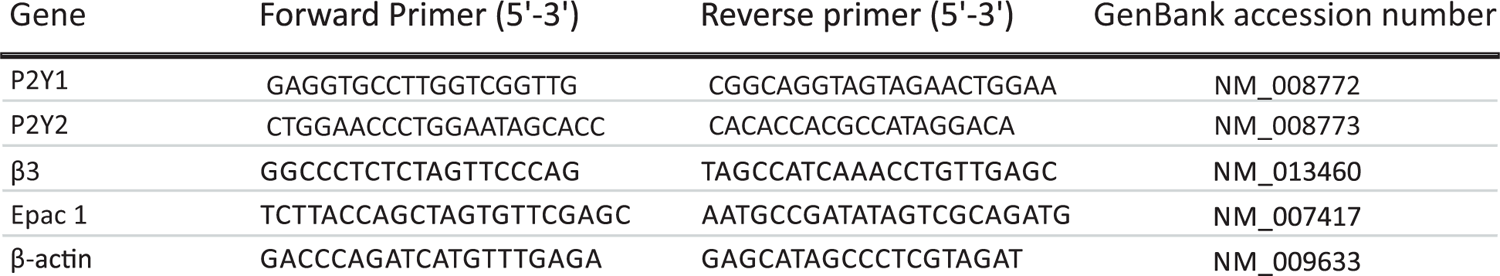

